# Cellular and Extracellular microRNA Dysregulation in LRRK2-Linked Parkinson’s Disease

**DOI:** 10.1101/2025.04.15.648997

**Authors:** Felix Knab, Jun-Hoe Lee, Raja Nirujogi, Kevin Menden, Fatma Busra Isik, Anto Praveen Rajkumar, Stefan Czemmel, Julia Fitzgerald, Thomas Gasser

## Abstract

**Background and objective:** The discovery of cell-free micro-RNAs in body fluids has made them a promising biomarker target in the field of neurodegenerative diseases. Although they have been reported to be differentially expressed in biofluids and tissues from sporadic Parkinson’s disease patients, it remains unclear whether similar observations can be made in patients with genetic forms of the disease and if miRNA profiles reflect mutation-specific pathogenic pathways. Since induced pluripotent stem cell-derived neurons represent a widely used research model for both sporadic and familial Parkinson’s disease, we sought to assess the usability of this model for the identification of differentially expressed cell-free micro-RNAs in the context of the Parkinson’s disease-related LRRK2 G2019S mutation in a proof-of-concept study.

**Materials and methods:** We isolated extracellular vesicles carrying cell-free RNA from patient-derived induced pluripotent stem cells carrying the LRRK2 G2019S mutation and their gene-corrected isogenic controls. After the generation of small-RNA libraries and differential expression analysis, we quantified expression levels of fourteen micro-RNAs in an independent batch of cell-free and cellular RNA via RT-qPCR. Finally, we quantified pRab10 levels as a proxy of LRRK2 activity and correlated observable changes to the miRNA expression levels.

**Results:** We successfully isolated extracellular vesicles from induced pluripotent stem cell-derived human dopaminergic neurons. We detected over 2000 different micro-RNAs of which 56 were differentially expressed. Dysregulation of four micro-RNAs was confirmed in an independent batch of cell-free RNA. We discovered a high correlation between changes in the cell-free and cellular micro-RNAomes. Finally, we showed poor correlation between LRRK2 expression or activity and miRNA expression levels.

**Conclusions:** Our results suggest that patients carrying the LRRK2 G2019S mutation display alterations in cellular and cell-free micro-RNA expression levels. Notably, the miRNA changes observed in this study did not follow a linear relationship with LRRK2 expression levels or kinase activity. Validation in larger cohorts will be necessary.

## Introduction

Parkinson’s disease (PD) is a neurodegenerative disorder with an estimated total of over six million affected patients worldwide [1]. Diagnosing PD, particularly in its early stages, remains a clinical challenge [2]. There is a great need for non-invasive markers identified in body fluids, as they could lay the foundation for a cost-effective and widely available indicator of clinical onset or disease progression.

Initially considered a sporadic disorder, PD is now known to have genetic underpinnings in 10– 15% of cases, displaying Mendelian inheritance patterns. Over the past few decades, more than ten genes have been implicated in familial Parkinson’s disease (fPD) [3]. Although these mutations account for only a small percentage of PD cases, they offer valuable insights into the disease’s complex and heterogeneous pathogenesis. Mutations in the Leucine-rich repeat kinase 2 (LRRK2) gene, such as the LRRK2 G2019S mutation, represent the most common genetic cause of fPD [4; 5] and due to reduced penetrance, LRRK2 mutations can also be found in patients with sPD (∼2%) [6]. This suggests that LRRK2 plays an important role in the pathogenesis of at least a specific subset of both sporadic and familial PD. Various functions of LRRK2 in cellular mechanisms related to Parkinson’s disease have been observed [7]. Often, the importance of the altered kinase activity of LRRK2 is highlighted. In many studies, the phosphorylated-to-total Rab10 ratios serve as a proxy for LRRK2 kinase activity [8; 9]. With LRRK2 inhibitors rapidly approaching clinical testing, a LRRK2-related non-invasive readout of target engagement or treatment efficacy would be of immense interest.

micro-RNAs (miRNA) have emerged as a promising target in the field of biomarker research [10]. miRNAs are small non-coding RNAs that post-transcriptionally regulate gene expression via degradation of mRNA or translational repression [11]. They are easily accessible as cell-free RNA in body fluids such as plasma or CSF [12] and are reported to be relatively stable [13]. While various mechanisms of active secretion into the extracellular space are being investigated [14; 15; 16], it seems clear that a relevant amount of cell-free RNA exists within extracellular vesicles (EVs) [17]. The term “extracellular vesicle” refers to membrane-enclosed vesicles with a size of 40 to 1000 nm that are secreted by cells and contain cytosolic components such as proteins and nucleic acids [18]. EVs have been intensively studied in past biomarker studies [19], as they can easily be isolated from biofluids such as plasma and CSF [20].

A vast number of both cellular and cell-free miRNAs are reported to be dysregulated in neurodegenerative diseases, including sPD [21] but very little information exists on miRNA dysregulations specific to fPD. It is known that mutated LRRK2 interferes with the cellular miRNAome [22; 23] and many miRNAs were predicted to target LRRK2 [24], though only a few have been experimentally validated.

In this proof-of-concept study, we wanted to identify differentially expressed cell-free miRNAs derived from neurons carrying the LRRK2 G2019S mutation compared to gene corrected isogenic controls. We thus differentiated human dopaminergic neurons (hDaNs) from two previously characterized induced pluripotent stem cell (iPSC) lines [25; 26] derived from patients with a LRRK2 G2019S mutation, used EVs as a source for cell-free RNA, and generated small-RNA libraries. We then compared miRNA expression from LRRK2 G2019S lines to isogenic gene-corrected control lines. We further wanted to know how well cell-free miRNA expression levels reflect the intracellular miRNAome. We therefor extracted cellular RNA from iPSC-derived hDaNs and again quantified miRNA expression levels. Finally, we tested whether changes of the observed miRNA profiles were related to LRRK2 expression levels or kinase activity.

## Materials and Methods

### Differentiation of Human Dopaminergic Neurons from Neuronal Progenitor Cells

Two lines of neural progenitor cells (NPCs) carrying the LRRK2 G2019S mutation or the gene-corrected LRRK2 wild type were previously generated from patient-derived iPSCs [25; 26]. Here, L1 and L2 refer to the two PD patients carrying the LRRK2 G2019S mutation from whom cells were derived. L1 Mut and L2 Mut denote the cell lines carrying the LRRK2 G2019S mutation, while L1 GC and L2 GC represent the gene-corrected isogenic control lines. When referring to both lines derived from a single patient (e.g., L1 GC and L1 Mut), we use the terms L1 lines or L2 lines, respectively. NPCs were cultured in base media (50% neurobasal (Thermo Fisher Scientific (Waltham, MA, USA), #21103-049), 50% DMEM/F12 (Thermo Fisher Scientific, #11-330-057), 1% penicillin/streptomycin (Merck (Darmstadt, Germany), #A2213), 1% GlutaMax (Thermo Fisher Scientific, #35050-038), 1% B27 supplement (without vitamin A; Thermo Fisher Scientific, #12587-010) and 0.5% N2 supplement (Thermo Fisher Scientific, #17502-048)) supplemented with 0.5 µM PMA (Merck, #540220), 3 µM CHIR (Axon Medchem (Groningen, Netherlands), #Axon1386) and 200 µM ascorbic acid (Sigma-Aldrich (St. Louis, MO, USA), #A4544). For each line, one cryotube of NPCs was thawed, and cells were split into three different 6-wells. The three wells of NPCs were expanded independently and treated as three differentiation replicates. Once NPCs reached 80% confluency, differentiation was started. For the differentiation into human dopaminergic mid-brain neurons (hDaNs), NPCs were incubated in base media supplemented with 20 ng/ml of BDNF (PeproTech (Hamburg, Germany), #450-02), 10 ng/ml of FGF8 (PeproTech, #100-25), 1 µM of PMA and 200 µM of ascorbic acid for 7 days (days 1 to 7). Next, cells were put in base media supplemented with 10 ng/ml of BDNF, 10 ng/ml of GDNF (PeproTech, #450-10), 1 ng/ml of TGF-β3 (PeproTech, #AF-100-36E), 200 µM ascorbic acid, 500 µM dbcAMP (PanReac AppliChem (Darmstadt, Germany), #A0455) and 10 µM DAPT (Selleckchem (Houston, TX, USA), #S2215) for the remainder of the differentiation (days 8 to 23). None of the used media contained fetal calve serum.

### Quantifying the Expression Levels of Neuronal Markers in hDaNs

To quantify the expression of neuronal and dopaminergic markers in the hDaNs, RT-qPCR was performed. Cellular RNA was isolated on day 23 of differentiation using the miRNeasy Tissue/Cells Advanced Mini Kit following the vendor’s instructions (QIAGEN (Hilden, Germany), #217604). The mRNA expression levels of the following genes were quantified: dopaminergic marker tyrosine hydroxylase (TH), mature neuron marker microtubule associated protein 2 (MAP2), midbrain marker forkhead box protein A2 (FOXA2), and LRRK2. RT-qPCR was carried out using the QuantiTect® SYBR®-Green RT-PCR Kit (QIAGEN, #204243, customized primer sequences are listed in Supplemental Table 1). GAPDH was used for internal normalization while RNA isolated from gene-corrected iPSCs derived from L2 was used as an external reference. RT-qPCR was performed on the LightCycler® 480 (Roche) and c(t)-values were extracted using the LightCycler® 480 software (version 1.5.1). The delta-delta-c(t) method was used to calculate the fold change (fc) expression levels of hDaNs over iPSCs.

### Immunocytochemistry of hDaNs

The hDaNs were split on day 8 of the differentiation process and seeded onto 12 mm-wide coverslips. Before seeding, cover slips were coated with 15µg/ml of Poly-DL-ornithine (Sigma-Aldrich, #P8638) at 37° C for 24 hours, followed by coating with 5µg/ml laminin (Sigma-Aldrich, #L2020) at 37° C for 4 hours. On day 23, cells were fixed by adding 4% PFA to the cells for 20 minutes. Cells were washed with PBS (Sigma-Aldrich, #D8537) three times before blocking, using PBS containing 10% normal goat serum (NGS) and 0.1% Triton-X for one hour. Blocking buffer was removed, and primary antibodies were added, resuspended in 5% NGS and 0.1% Triton-X in PBS. MAP2 antibody was used in a 1:2000 dilution (abcam (Cambridge, UK), #ab5392) and Tyrosine Hydroxylase (TH) antibody was added in a 1:1000 dilution (Pel-Freez (Rogers, AR, USA), #P40101-150). Primary antibodies were incubated overnight at 4° C before cells were washed three times with PBS. Secondary antibodies were added in 5% NGS and 0.1% Triton-X. Anti-Chicken AlexaFluor-647 (Thermo Fischer Scientific, #A21449) and Anti-Rabbit AlexaFluor-488 (Thermo Fischer Scientific, #A11070) were used in a dilution of 1:1000. After one hour of incubation at 4° C, cells were washed three times, and nuclei were stained with Hoechst (Molecular Devices (Urstein, Austria), #H3569) for five minutes. Finally, coverslips were mounted using Dako mounting medium (Aligent (Santa Clara, CA, USA), #S3023). Ten positions were imaged per differentiation and line on a Zeiss Imager.Z1 using the ZEN software (blue edition, Zeiss, (Oberkochen, Germany)). Prior to analysis, Z-stacks were projected, and brightness was adjusted for each channel. The total number of intact nuclei was counted, and percentage of TH and MAP2-positive cells was calculated.

### Isolation of Extracellular Vesicles from Cell Culture Media

Starting on day 14 of the differentiation protocol, conditioned cell culture media (CCM) was collected from hDaNs every three days in order to isolate extracellular vesicles. The CCM was centrifuged at 300g for ten minutes to deplete cell debris before the supernatant was filtered through a 0.22 µm Steriflip filter (Merck, #SCGP00525). The CCM was then concentrated to 1 ml using Amicon® Ultracel Centrifugal filters (Merck, #UFC901024) and centrifugation at 2000g at 4° C for 30 minutes. Next, the concentrate was recovered and transferred to a 1.5-ml tube. Subsequently, 500 µl of Total Exosome Isolation Reagent (Thermo Fischer Scientific, #4478359) was added to 1 ml of concentrated CCM. The samples were then incubated overnight and centrifuged at 10 000g for one hour. The supernatant was discarded, and tubes were centrifuged for 5 minutes at 10 000g. Next, pellets were washed three times with 1 ml PBS before resuspension in 200ul of PBS containing cOmplete protease inhibitor (Sigma-Aldrich, #11873580001) and phosphatase inhibitor (Sigma-Aldrich, #4906837001).

### Western Blotting of LRRK2 and Extracellular Vesicle Markers

For cell protein extracts, hDaNs from a 6-well were lysed on day 23 of differentiation in 200 µl ice-cold lysis buffer (1% Triton X-100 in PBS) and incubated on ice for five minutes. Resulting cell suspensions were consequently centrifuged at 13 000 g at 4°C for 15 minutes. As for Western blot analysis, 25 to 75 µg of the cell lysates were mixed with 2x Laemmli sample buffer (Sigma-Aldrich, #S3401-10VL), separated in an 8-10% polyacrylamide gel, and transferred to a 0.45 µm Immobilon-P PVDF membrane (Merck, # IPVH00010). The membranes were blocked in 5% (w/v) milk (PanReac AppliChem, #A0830) in TBS-T for 1h at RT, incubated with anti-LRRK2 (dilution: 1:500, abcam, #ab133474) or anti-ß-actin (1:5000, abcam, #ab6276) antibodies overnight at 4°C and with secondary HRP-conjugated antibody (anti-rabbit: 1:5000, Cell Signalling (Cambridge, UK), #7074; anti-mouse: 1:10 000, Bio-Rad (Hercules, California, USA), #172-1011) for 1h at RT. For detection, the membranes were incubated with ECL prime western blot detection reagents (Merck, #GERPN2232) and the signal was detected using Amersham Hyperfilm ECL (Thermo Fisher Scientific, # 10534205). For quantification, the chemiluminescence signal intensity was measured using ImageJ software, and LRRK2 protein levels were normalized to levels of ß-actin expression. For the detection of EV markers, slight modifications were made. Briefly, EV samples were incubated in SDS sample buffer (0.375 M Tris-Cl, 12% SDS, 60% Glycerol, 0.6 M DTT, pH 6.8) for 10 minutes at 95° C, run on a 10% polyacrylamide gel, and transferred onto a nitrocellulose membrane using the iblot 2 Dry Blotting System. Anti-Alix (1:250, NovusBio (Littleton, Colorado, USA), #NBP1-90201), anti-Flotillin-1 (1:1000, Cell Signalling, #18634) and anti-CD81 (1:1000, Santa Cruz (Dallas, Texas, USA), sc-166029) antibodies were used to detect corresponding EV-specific markers. Anti-GM130 antibody (1:1000, Cell Signaling, #12480) was used as a negative control. As a positive detection control, 10 µg of neuronal cell lysates were used.

### Nanoparticle Tracking Analysis

Nanoparticle Tracking Analysis (NTA) was used to measure particle size and concentration. The Nanosight NS300 and NanoSight NTA 3.0 0068 software from Malvern Panalytical in Kassel, Germany, were used. For optimal performance, samples were diluted 1:100 to 1:500 in PBS prior to measurement. Each sample was recorded and measured five times.

### Cryo Transmission Electron Microscopy

Cryo Transmission Electron Microscopy (TEM) of the EV samples was conducted in the Nanoscale and Microscale Research Centre (nmRC) of the University of Nottingham, UK. The nmRC has previously published its protocol for preparing EV samples for cryo-TEM, and the protocol was adapted for this manuscript [27]. Holey carbon TEM grids (EM resolutions, Sheffield, UK, #HC300Cu) were used. The EV samples were left to adsorb onto the grids (5 µL/grid) for two minutes, then excess solution was removed using filters. After blotting the EV samples for a second, they were frozen in liquid ethane using a Gatan CP3 plunge freezing unit (Ametek, Leicester, UK). Frozen samples were loaded onto a FEI Tecnai G2 12 Bio-twin TEM, and Cryo-TEM was completed with an accelerating voltage of 100 kV. Images were obtained using an in-built Gatan SIS Megaview-IV digital camera.

### Generation of Small-RNA Libraries using Cell-free RNA from hDaNs

For the generation of small-RNA libraries, RNA was isolated directly from EVs derived from supernatant from either day 14 to 23 (L1 lines) or day 14 to 17 (L2 lines). RNA was isolated using the miRNeasy Serum/Plasma Advanced Kit (QIAGEN, #217204) and following the vendor’s instructions. After isolation, RNA concentrations were measured using the QubitTM microRNA Assay Kit (Thermo Fischer Scientific, #Q32880) and electropherograms were generated using the high-sensitivity RNA ScreenTape® (Agilent, Santa Clara, California, USA, #5067-5579). For the generation of the small RNA libraries, the SMARTer® smRNA-Seq Kit for Illumina® (Takara (Tokyo, Japan), #635030) was used, and the vendor’s instructions were followed with slight modifications. Briefly, 5 ng of RNA were used for the initial polyadenylation step. This was followed by cDNA synthesis using PrimeScript RT. Finally, cDNA was used for amplification via PCR using one forward primer and differently barcoded reverse primers. The libraries were then bead-purified by first incubating AMPURE beads (Beckmann Coulter (Brea, California, USA), #A63881) with the samples at room temperature for five minutes in a sample-to-bead ratio of 1:1.8. The beads were then incubated on a magnetic rack for another three minutes. The supernatant was discarded, and the beads were washed twice with 80% ethanol and dried for five minutes. Next, the beads were resuspended in 32 µl of water and incubated at room temperature for five minutes. Afterwards, the tubes were placed onto the magnetic rack, and the supernatant was transferred to a new tube. For size exclusion, the same protocol was repeated using SPRI beads (Beckmann Coulter, #B23318) and a sample-to-beads ratio of 1:1. For the generation of electropherograms of the libraries, the high-sensitivity D1000 ScreenTape® (Agilent, # 5067-5584) was used. Finally, libraries were sequenced on a Nextseq550 mid-output, 150 cycles v2.5 flowcell spiking in 30% PhiX control v.3. Although the analysis of mRNA is not part of the present study, sequencing was done at a higher length than usual for small RNA sequencing to allow the inclusion and better mapping of potentially present mRNAs.

### Differential Expression Analysis of Library Sequencing Data

The raw sequence data was transferred to the Quantitative Biology Centre (QBiC), Tübingen, Germany, for geo-redundant long-term storage, data management, and bioinformatics analysis. For data processing, the Nextflow-based nf-core/smrnaseq pipeline (v. 1.2.0) development branch was forked (https://github.com/qbic-projects/smrnaseq, commit: 306b024) to allow for the input of custom adapter trimming parameters. The nf-core/smrnaseq includes various bioinformatic tools, such as FastQC (v0.11.9) (Andrews, 2010) to determine the quality of the FASTQ files. This was followed by read length selection (maximum 40 bp) and adapter trimming using Trim Galore (v0.6.h) [28], based on the recommended parameters by the Takara SMARTer® smRNA-Seq Kit. Subsequently, the filtered reads were mapped to the miRBase [29] database of mature and hairpin human miRNAs using Bowtie1 (v. 1.3.0) [30]. The miRNA-seq data quality was assessed using mirTrace (v1.0.1) [31], followed by an aggregation of the quality control of the analyses with MultiQC (version 1.7; http://multiqc.info/) [32]. Differential expression analysis on the miRNA sequences was then performed using the read counts mapped by Bowtie1. The analysis was performed in R (v3.5.1) using DESeq2 (v1.22.1) [33], through a fork of the Nextflow-based qbic-pipelines/rnadeseq pipeline (v1.32). This fork (https://github.com/qbic-projects/QSCNN, commit: c1aded5) was created to account for the lower number of expressed miRNAs (hairpin, mature) in comparison to typical gene expression data; hence, the variance stabilizing transformation parameter (default: 1000) was adjusted to 400. The linear model employed to model gene expression in DESeq2 was "∼ Patient + condition_genotype" to account for differences in patients while checking for the effect of the main experimental factor "condition_genotype" (G2019S (mutation in LRRK2) vs. gene-corrected control). The miRNA sequences were considered differentially expressed when the Benjamini-Hochberg multiple testing adjusted p-value [34] was smaller than 0.05 (padj < 0.05). Multiple testing corrections were used to minimize the number of false positives.

### Relative Quantification of miRNAs in Cellular and Cell-free RNA from hDaNs via RT-qPCR

For validation of the differentially expressed miRNAs found in the libraries, a new batch of cell culture-derived EVs was generated, and cell-free RNA was isolated as described previously. For optimal comparison, EVs were isolated from supernatant collected from days 14 through 23 for both the L1 and L2 lines. Cellular RNA was isolated from hDaNs on day 23 of the differentiation. For quantification of specific miRNAs, RT-qPCR was performed using the miRCURY LNA SYBR Green PCR Kit (QIAGEN, #339345) and the vendor’s instructions were followed with slight modifications. Briefly, for reverse transcription, 2 µl of 5x miRCURY RT Reaction Buffer were mixed with 1 µl of 10x RT Enzyme Mix and 7 µl of cell-free RNA or 10 ng of cellular RNA. The sample was incubated for one hour at 42 °C, followed by an inactivation step at 95 °C for 5 minutes. Cell-free cDNA was diluted at 1:30, while cellular cDNA was diluted at 1:60 using nuclease-free water. For the RT-qPCR reaction, 5 µl of 2x miRCURY SYBR Green Master Mix was added to 1 µl of miRCURY LNA miRNA PCR Assay (QIAGEN, #339306) and 4 µl of diluted cDNA template. Three technical replicates per sample and target were measured. Samples were analyzed in a LightCycler® 480 after an initial PCR heat activation step for 2 minutes at 95°, followed by 2-step cycling. After 45 runs, melting curve analysis was performed using the LightCycler® 480 Software version 1.5.1, and targets with unspecific amplifications (c(t) ≥40) were excluded. For the extraction of c(t)-values, LinRegPCR version 2021.2 was used [35]. The delta-delta-c(t) method was used to calculate the relative expression of miRNAs and miRNA-16 was used as an internal reference. miRNAs with a fc of ≥1.5 or ≤0.5 were considered differentially expressed. A miRNA was considered validated if direction of dysregulation matched the observation in the library.

### Quantification of phosphorylated Rab10 in Cell Lysates and Extracellular Vesicles

Cell lysates as well as EV samples were reduced and alkylated by adding 5mM DTT and incubated at 56°C for 30 minutes followed by alkylated by adding 20mM Iodoacetamide and incubated in dark for 30 min. followed by samples were resolved on a NuPage 10% Bis-Tris gel (ThermoFischer Scientific, NP0301BOX) Bands containing proteins with a size of 20-30 kDa were excised for In-gel digestion [36]. Briefly, bands were cut into 1mm gel-pieces and then destained by adding 250µL of destaining buffer (40mM Ammonium bicarbonate in 40% (vol/vol) Acetonitrile in milli-Q water) and incubated on a Thermomixer for 20 minutes with an agitation at 1200 rpm at room temperature. Following the buffer was discarded and repeated this step again to ensure complete destaining. Further, gel pieces were dehydrated by adding 250µL of 100% (vol/vol) Acetonitrile and incubated on a Thermomixer for 20 minutes with an agitation at 1200 rpm at room temperature, this step was repeated again until the gel-pieces were completely dehydrated. 500ng of Trypsin was added in a 200µL of trypsin buffer (0.5% (wt/vol) sodium deoxycholate in 20mM TEABC pH 8.0 in milli-Q water). The tryptic digestion was continued for overnight by placing the samples on a Thermomixer at 37oC by agitating at 1200 rpm. Peptide extraction was performed by adding 100 µL of 99% Isopropanol (vol/vol) in 1% TFA and incubated on a Thermomixer at room temperature by agitation at 1200 rpm for 20 minutes. The supernatant was transferred to new 1.5 ml protein lo-binding tube and the extraction was continued another two times, the eluates were pooled and directly loaded on Stage-tips for SDB-RP clean up and the eluted peptides were subjected to vacuum dryness on a SpeedVac concentrator. The peptides were then stored in -80 freezer until LC-MS/MS analysis. For this, peptides were dissolved in LC-buffer (3% ACN vol/vol in 0.1% formic acid vol/vol in milli-Q water) and was spiked with an equimolar ratio of 25 fmol heavy peptide mix containing heavy pRab10 (FHTITp**T**SYYR*), heavy non-phospho Rab10 (FHTITTSYYR*) and two total Rab10 peptides (NIDEHANEDVER* and AFLTLAEDILR*). Peptide mixture was then loaded on Evotips for targeted mass spectrometry analysis as described in PMID: 34125248. The data was acquired in PRM mode on Orbitrap Exploris 240 mass spectrometer interfaced with EvoSep LC system. The mass spectrometry data was processed using Skyline software suite for peak picking and the ratio of light/heavy for pRab10, non-phospho Rab10 was used in measuring stoichiometry of phosphorylation [36; 37]. Finally, pRab10/Rab10 ratios were normalized to the LRRK2/ß-Actin ratio and values were logarithmized.

### Data analysis

Statistical analysis, except for the analysis of the small RNA-Seq data, was performed using either GraphPad Prism software, version 9.3.0. (La Jolla, CA, United States) or R-Studio, version 4.3.0. Normal distribution of the data was checked using QQ-plots. Non-parametric data (mRNA expression levels) was log-transformed, the alpha level was set at 0.05, and mean differences and standard deviations are reported. Ordinary Two-Way-ANOVA followed by Šidák’s test was used to test for differences between the genotypes when analyzing the mRNA expression levels and the results of the immunofluorescence staining. Factors were mutation status (LRRK2 G2019S and LRRK2 wildtype) and target (ICC: MAP2 and TH; mRNA expression: MAP2, TH, FOXA2, LRRK2), L1 and L2 lines were tested separately. Two-Way-ANOVA followed by Tukey’s post hoc test was used to test for differences of LRRK2 expression and pRab10 ratios in cell lysates and EVs comparing the two genotypes. Factors were mutation status (LRRK2 G2019S and LRRK2 wildtype) and patient line (L1 and L2). Linear regression models were applied to assess the association between log10-transformed miRNA expression values and either log2-transformed LRRK2/ß-actin ratios or the pRab10/total Rab10 to LRRK2/ß-actin ratio. P-values and R²-values are given. Individual differentiations are indicated as n_Diff_, technical replicates in qPCR experiments as n_Tec_ and the number of excluded miRNA targets as n_Exc_. Figures were built using either the GraphPad Prism software or RStudio (version 2022.12.0+353) [38].

## Results

### Immunocytochemistry of Neuronal and Dopaminergic Marker

Immunofluorescence staining of hDaNs revealed the neuronal morphologies of the cells and the presence of the dopaminergic marker TH and the mature neuron marker MAP2 (Figures 1A and 1B). In L1 GC, 73.6% (SD: ±5.8%) of the cells were MAP2 positive, compared to 75.1% (SD: ±11.6%) in L1 Mut. Furthermore, 29.6% (SD: ±2.2 %) of the cells in L1 GC and 41.6% (SD: ±2.8%) of the cells in L1 Mut showed TH expression (Figure 1C). Ordinary Two-Way-ANOVA was performed to test for differences in percentages of TH or MAP2 positive cells between the two genotypes, and no significant interaction was observed between target and genotype (F(1, 8) = 1.81, p = 0.215). Simple main effects analysis showed that target (F(1, 8) = 99.72, p<0.0001) had a significant effect on the percentage of positively expressing cells but not genotype (F(1, 8) = 2.971, p = 0.123). After correction for multiple comparison using Šidák’s test, neither the difference of TH positive cells (mean difference: 0.12 percentage points (p.p.), q = 3.07, DF = 8, p = 0.211) nor of MAP2 positive cells (mean difference: 0.01 p.p., q = 0.377, DF = 8, p = 0.993) was observed to be significantly different between genotypes.

**Figure 1.**
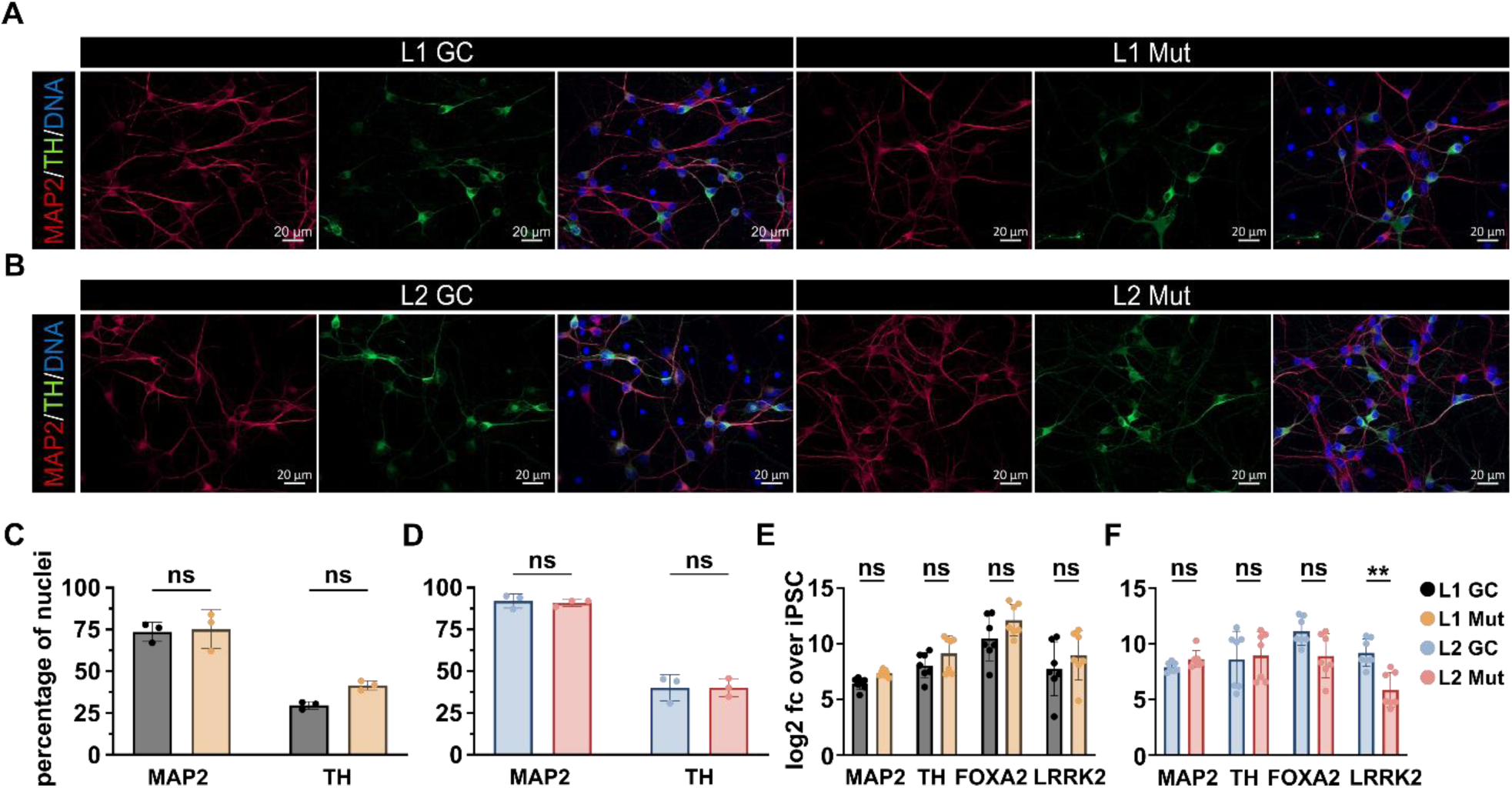
Validation of neuronal identity**. (A)** Representative images of immunofluorescence staining of hDaNs from the L1 lines and **(B)** the L2 lines on day 23 of the differentiation. Dopaminergic marker TH was stained together with neuronal marker MAP2 and DNA. **(C)** Percentages of MAP2+ and TH+ cells were calculated for both the L1 and **(D)** the L2 lines. 10 different positions were analyzed (n_Diff_ = 3). Error bars indicate standard deviation. **(E)** RT-qPCR was performed on cellular RNA isolated on day 23 from the L1 and **(F)** the L2 lines to quantify expression of MAP2, TH, FOXA2 and LRRK2 on mRNA level (n_Diff_ = 7). Results are shown as log2 fc of the expression in iPSC control samples. Error bars indicate standard deviation.

The number of MAP2-positive cells was observed to be 92.1% (SD: ±4.2%) in L2 GC and 90.8% (SD: ± 2.0%) in L2 Mut. On the other hand, the number of cells showing positive TH expression was 40.0% (SD: ±8.0%) in L2 GC and 40.2% (SD: ±5.5%) in L2 Mut (Figure 1D). Ordinary Two-Way-ANOVA again showed no significant interaction between target and genotype (F(1, 8) = 0.050, p = 0.829). Simple main effects analysis showed that target (F(1, 8) = 271.9, p<0.0001) had a significant effect on the percentage of positively expressing cells but not genotype (F(1, 8) = 0.030, p = 0.867). Importantly, no significant difference in TH positive cells (mean difference: <0.01 p.p., q = 0.049, DF = 8, p>0.999) or MAP2 positive cells (mean difference: 0.01 p.p., q = 0.396, DF = 8, p = 0.992) was found between genotypes.

### mRNA Expression Levels of Neuronal and Dopaminergic Marker

Our quantification of TH, MAP2, and FOXA2 mRNA in all four differentiated lines showed strong upregulation compared to iPSCs. (Figures 1E and 1F). As for LRRK2, Two-Way-ANOVA indicated that there was no statistically significant interaction between the effects of genotype and target in the L1 lines ((F3, 48) = 0.120, p = 0.948). Simple main effects analysis showed that both genotype and target had a statistically significant effect on the expression levels (target: F(3, 48) = 17.90, p < 0.0001; genotype: F(1, 48) = 7.926, p = 0.007). After correction for multiple comparisons using Šidák’s test, no significant difference between L1 Mut and L1 GC was found for any of the gene expression levels. In contrast, ordinary Two-Way-ANOVA in the L2 lines indicated a significant interaction between genotype and target (F3, 48) = 5.168, p = 0.004). Simple main effects analysis showed that both genotype and target had a statistically significant effect on the expression levels (target: F(3, 48) = 5.817, p = 0.002; genotype: F(1, 48) = 6.604, p = 0.013). After correction for multiple comparisons using Šidák’s test, the levels of LRRK2 expression were found to be significantly decreased in L2 Mut compared to L2 GC (t = 3.839, DF = 48, p = 0.0014).

### Confirmation of Presence of Extracellular Vesicles

After validating the neural identity of the cells, we wanted to confirm the successful isolation of EVs from cell culture supernatants. In Cryo-TEM, particles appeared as solitary, spherical, and membrane-encapsulated structures, as previously reported [39] (Figure 2A). The size of particles from all four lines ranged from 30 nm to 200 nm (measured using NTA, Figure 2B). The mean size of particles isolated from L1 GC was 80.53 nm (SD: 6.89, n_Diff_ = 6) and 77.86 nm (SD: 5.05, n_Diff_ = 6) in L1 Mut. The average size of particles in the L2 GC was 81.08 nm (SD: 7.54, n_Diff_ = 6) and 80.33 nm (SD: 9.01, n_Diff_ = 6) in L2 Mut. Furthermore, we tested for the presence of EV protein markers. The cytoplasmic protein Alix and the membrane-bound proteins Flotillin-1 and CD81 were detected in western blots, while GM130 was barely detectable [40] (Figure 2C). Taken together, particle size, the presence of EV protein markers and Cryo-TEM images confirmed the presence of EVs, following previously published guidelines [41].

**Figure 2.**
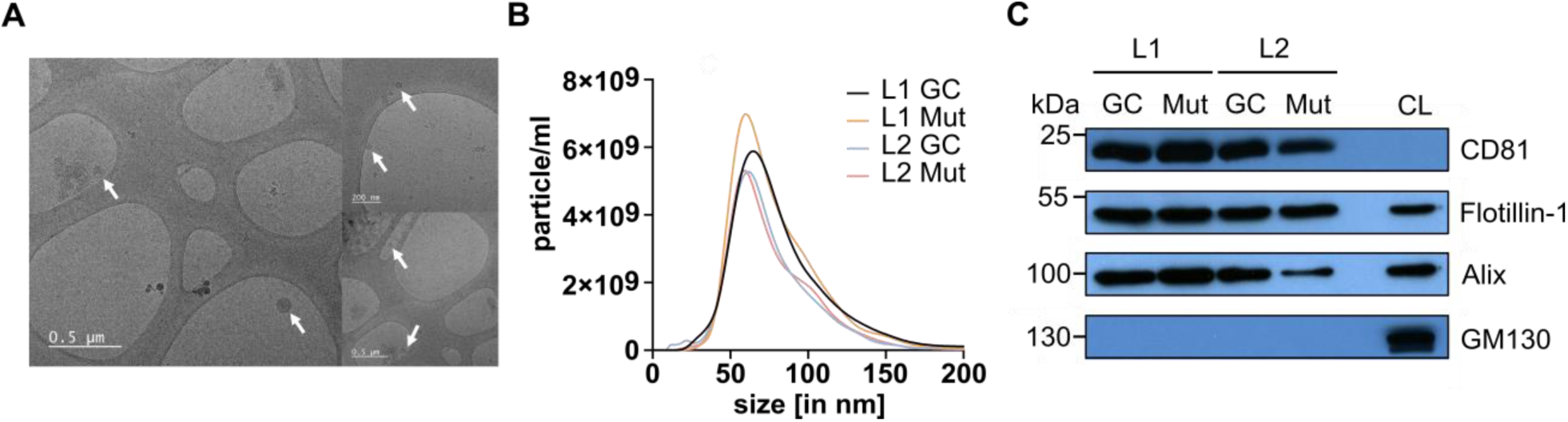
Characterization of neuronal EVs. **(A)** Representative Cryo-EM images of spherical and membrane-encapsulated vesicles. **(B)** NTA measurements of EVs derived from all four lines. Size ranged between 30 to 200 nm. **(C)** Western blotting of CD81, Flotillin-1 and Alix revealed presence of vesicle markers in the EV samples. The Golgi apparatus associated protein GM130 was used as a negative control. Neuronal cell lysates (CL) were used as a positive control.

### Quality Control of small RNA Libraries

Notably, RNA input from exosomes did not contain typical rRNA peaks that are usually used to verify RNA integrity (Supplemental Figure 1). This has previously been reported to be the case for exosomal RNA[14]. The mean GC percentage of the reads was 55.17% (SD: ±4.47), while the mean percentage of duplicate reads was 53.39% (SD: ±9.71). The mean percentage of miRNAs present in the libraries was 2.36% (SD: ±1.71). The percentage of reads that are mapped to miRBASE is 1.78 % (SD: ±1.01) for mature RNAs and 3.38 % (SD: ±1.75) for hairpin RNAs. These quality control values and statistics are generally in line with previously published small RNA libraries [42]. Further details (including the proportion of biotypes) are reported in Supplemental Figure 2 and Supplemental Tables 2-5.

### Small-RNA Libraries Reveal a Subset of Differentially Expressed miRNAs

A total of 2611 and 2608 miRNAs were detected in the generated libraries from the L1 and L2 lines, respectively. Both libraries shared a common set of 2091 (67%) miRNAs (Supplemental Figure 3A). 798 and 910 miRNAs did not pass our fc threshold, respectively (Supplemental Figure 3B). After correction for multiple testing, 19 miRNAs were significantly upregulated and 12 downregulated in L1 Mut compared to L1 GC (Figures 3A and 3C, Supplemental Table 6). In the L2 Mut, 11 miRNAs were significantly upregulated and 14 were significantly downregulated (Figures 3B and 3D, Supplemental Table 7). Despite some overlap of up- or downregulated miRNAs before correction for multiple testing between the two lines, they did not share any significantly dysregulated miRNAs after correction (Supplemental Figures 5C and 5D). When applying a less stringent p-value threshold of p≤0.1, miR-8064 was identified as significantly dysregulated in both libraries.

**Figure 3.**
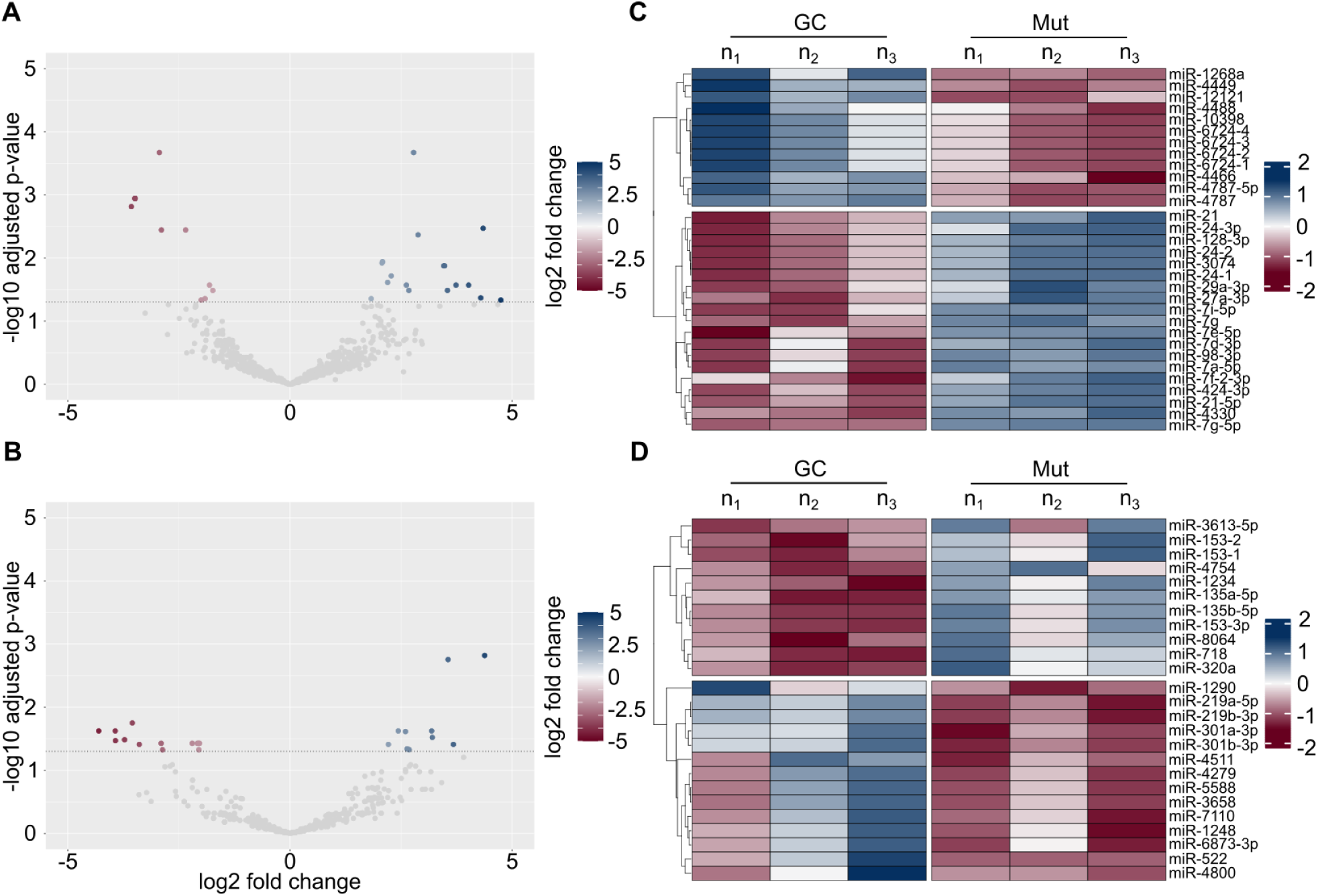
Results of the small-RNA-libraries generated from cell-free RNA of hDaNs**. (A)** A total of 2611 miRNAs were detected in the L1 libraries. 31 miRNAs passed the threshold and were considered differentially expressed (red dots). Log2 fc is expressed as change of L1 Mut over L1 GC. **(B)** In the L2 libraries, a total of 2608 miRNAs were found. 25 miRNAs were differentially expressed in L2 Mut compared to L2 GC (red dots). **(C)** vst-transformed gene counts were normalized from 0 to 100 for each target. In L1 Mut, of the 31 miRNAs that were differentially expressed, 19 were upregulated and 12 were downregulated. **(D)** In L2 Mut, out of the 25 differentially expressed miRNAs, 11 miRNAs were upregulated and 14 were downregulated.

### Validation of Differentially Expressed miRNAs

Next, after assessing the variation between technical replicates in the libraries for each miRNA, 14 miRNAs with small technical variability were selected for validation via RT-qPCR (Supplemental Table 8). A new and independent batch of EVs was generated and cell-free RNA was isolated and used for this validation step. Two miRNAs (miR-718 and miR-1234) were excluded due to unspecific amplification as indicated by the RT-qPCR melting curve. Of the tested miRNAs, two miRNAs could be validated in the L1 lines (log2 fc value: let-7g-5p: 1.99; miR-21-5p: 1.84), and the L2 lines (log2 fc: miR-135a-5p: 0.87, 153- 3p: 2.45) (Figure 4A). miR-135a-5p, identified from the L2 library, was also upregulated in cell-free RNA derived from L1 (log2 fc: 1.18). Table 1 summarizes the results from the RT-qPCR experiment using cell-free RNA.

**Figure 4.**
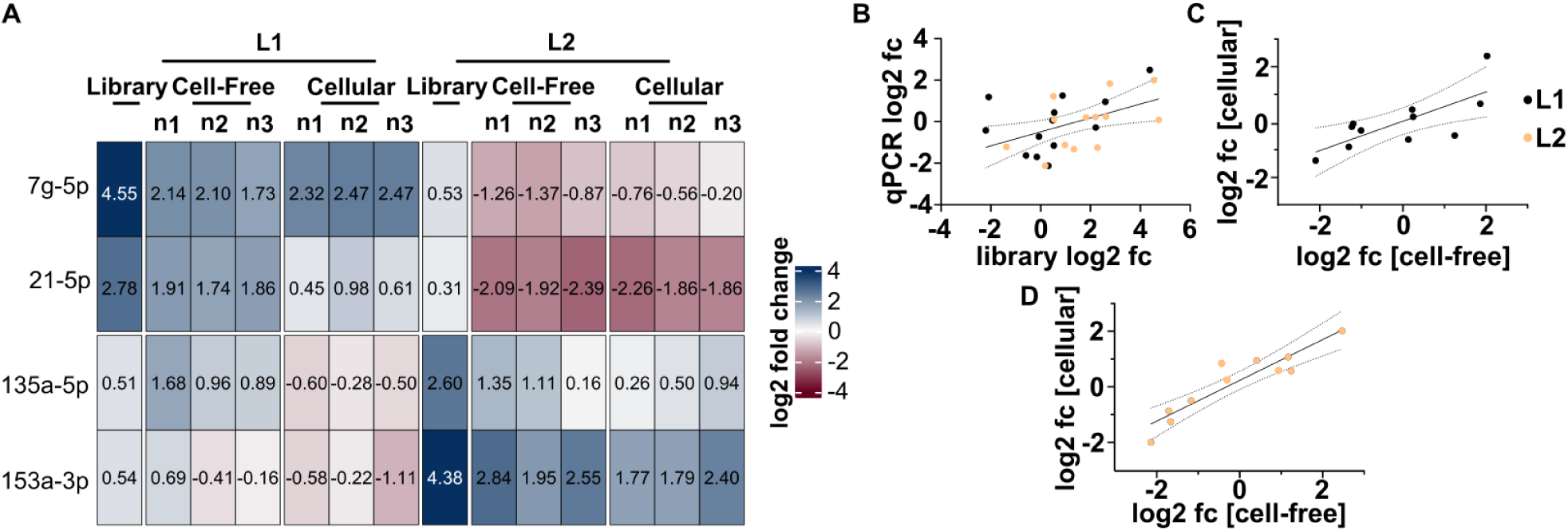
Results of the validation RT-qPCR using cell-free and cellular RNA. **(A)** The heatmap illustrates log2 fold-change values obtained from both library and RT-qPCR experiments for those miRNAs, that could be validated. RT-qPCR was conducted using both cell-free and cellular RNA. In L1, the dysregulation of let-7g-5p and miR-21-5p was successfully validated, while in L2, the dysregulation of miR-135a-5p and miR-153a-3p was confirmed. **(B)** Pearson’s correlation analysis revealed a statistically significant correlation between miRNA expression levels measured in cell-free RNA libraries and RT-qPCR, supporting the reliability of our findings. (**C**) Additionally, we conducted a correlation analysis of the log2 fold changes in miRNA expression between cellular and cell-free RNA in both L1 and (**D**) L2. This analysis included data from miRNAs whose expression changes observed in the libraries could not be independently validated. We identified a significant correlation between the two compartments, indicating that alterations in cell-free miRNA expression reflect those in the cellular miRNAome.

**Table 1.** Overview of the validation RT-qPCRs in cell-free RNA. Targets where dysregulation crossed the fc threshold of either ≥1.5 or ≤0.5 over the gene corrected control are highlighted in bold. A * highlights those miRNAs, where dysregulation found in the library was confirmed.

| target | fc (L1) | fc (L2) |
| --- | --- | --- |
| miR-21-5p* | **3.58 (SD ± 0.21)** | **0.23 (SD ± 0.04)** |
| let-7g-5p* | **4.00 (SD ± 0.59)** | **0.45 (SD ± 0.08)** |
| miR-128-3p | **0.42 (SD ± 0.07)** | 1.35 (SD ± 0.17) |
| miR-29a-3p | 1.19 (SD ± 0.14) | 1.04 (SD ± 0.15) |
| miR-27a-3p | 1.17 (SD ± 0.24) | **0.31 (SD ± 0.01)** |
| miR-24-3p | 1.15 (SD ± 0.28) | **0.32 (SD ± 0.02)** |
| miR-424-3p | 1.06 (SD ± 0.22) | 0.61 (SD ± 0.09) |
| miR-135a-5p* | **2.34 (SD ± 0.76)** | **1.94 (SD ± 0.73)** |
| miR-219a-5p | **0.23 (SD ± 0.04)** | **2.28 (SD ± 0.57)** |
| miR-301a-3p | **0.40 (SD ± 0.02)** | 0.75 (SD ± 0.10) |
| miR-320a | **0.43 (SD ± 0.02)** | 0.82 (SD ± 0.04) |
| miR-153a-3p* | 1.08 (SD ± 0.46) | **5.62 (SD ± 1.67)** |

Next, the respective log2 fc values from the libraries were plotted against the log2 fc from the RT-qPCR experiments to further assess the reliability of our findings. Using Pearson’s correlation, we found a statistically significant correlation between the values measured in the libraries and via RT-qPCR (r(24) = 0.48, p = 0.014, R^2^ = 0.23) (Figure 4B).

### Differentially Expressed Cell-Free RNAs Are Indicative of Changes in the Cellular miRNAome

We proceeded to quantify the expression levels of the 14 miRNAs selected from our libraries using cellular RNA (Figure 4A). Interestingly, the direction of dysregulation was mostly identical between cellular and cell-free RNA. Results are summarized in Table 2. To further analyze the relation between cellular and cell-free RNA, we performed correlation analysis on their respective log2 transformed fc values. In both L1 Mut (r(9) = 0.738, p = 0.0095, R^2^ = 0.545) and L2 Mut (r(9) = 0.922, p < 0.0001, R^2^ = 0.851), the correlation was statistically significant (Figures 4C and 4D). This indicates that the differential expression levels observed in cell-free miRNA are indicative of changes in the cellular miRNAome.

**Table 2.**
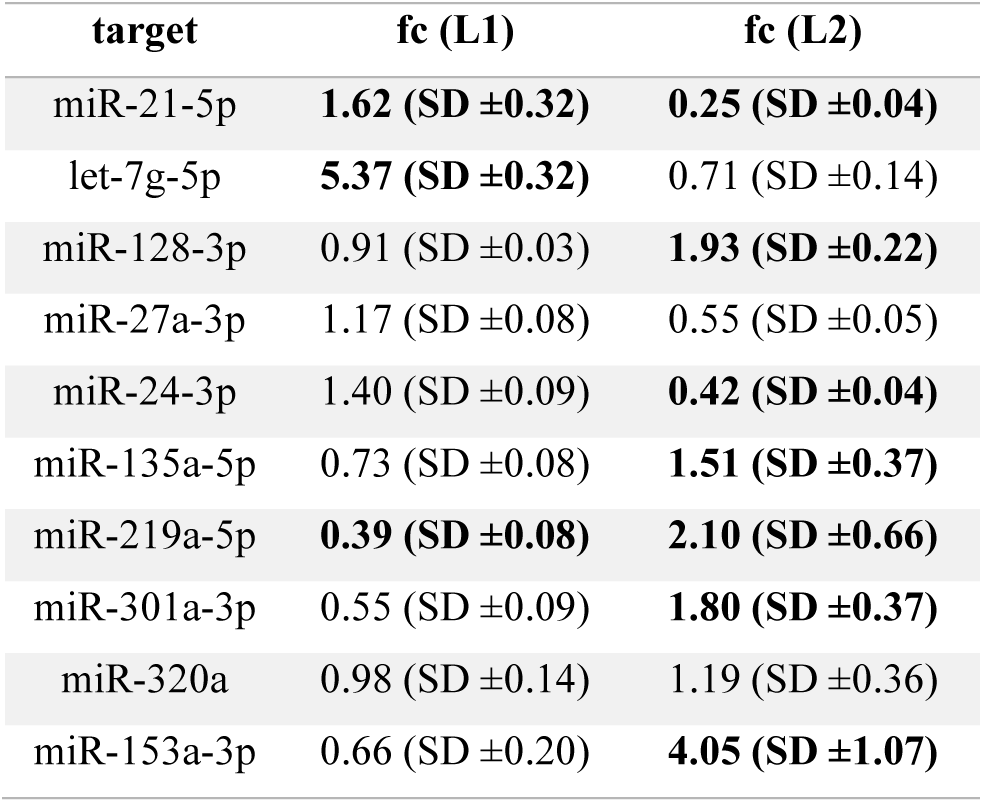
Overview of the RT-qPCRs results using cellular RNA. Targets where dysregulation crossed the threshold of either ≥1.5 or ≤0.5 over the gene corrected control are highlighted in bold.

| target | fc (L1) | fc (L2) |
| --- | --- | --- |
| miR-21-5p | **1.62 (SD $\pm 0.32$)** | **0.25 (SD $\pm 0.04$)** |
| let-7g-5p | **5.37 (SD $\pm 0.32$)** | 0.71 (SD $\pm 0.14$) |
| miR-128-3p | 0.91 (SD $\pm 0.03$) | **1.93 (SD $\pm 0.22$)** |
| miR-27a-3p | 1.17 (SD $\pm 0.08$) | 0.55 (SD $\pm 0.05$) |
| miR-24-3p | 1.40 (SD $\pm 0.09$) | **0.42 (SD $\pm 0.04$)** |
| miR-135a-5p | 0.73 (SD $\pm 0.08$) | **1.51 (SD $\pm 0.37$)** |
| miR-219a-5p | **0.39 (SD $\pm 0.08$)** | **2.10 (SD $\pm 0.66$)** |
| miR-301a-3p | 0.55 (SD $\pm 0.09$) | **1.80 (SD $\pm 0.37$)** |
| miR-320a | 0.98 (SD $\pm 0.14$) | 1.19 (SD $\pm 0.36$) |
| miR-153a-3p | 0.66 (SD $\pm 0.20$) | **4.05 (SD $\pm 1.07$)** |

### miRNA Expression Changes Show No Linear Relationship with LRRK2 Protein Levels or Activity

To further investigate the differences observed in LRRK2 mRNA expression levels and miRNA profiles across the four cell lines, we next assessed LRRK2 protein expression via Western blotting (Figure 5A). In the L1 cell lines, L1 GC exhibited a significantly lower LRRK2 protein expression compared to L1 Mut (U = 0, n = 6, p = 0.002). Conversely, in the L2 lines, LRRK2 protein levels were markedly higher in L2 GC compared to L2 Mut (U = 0, n = 6, p = 0.002), consistent with the corresponding mRNA expression data (Figure 5B).

**Figure 5.**
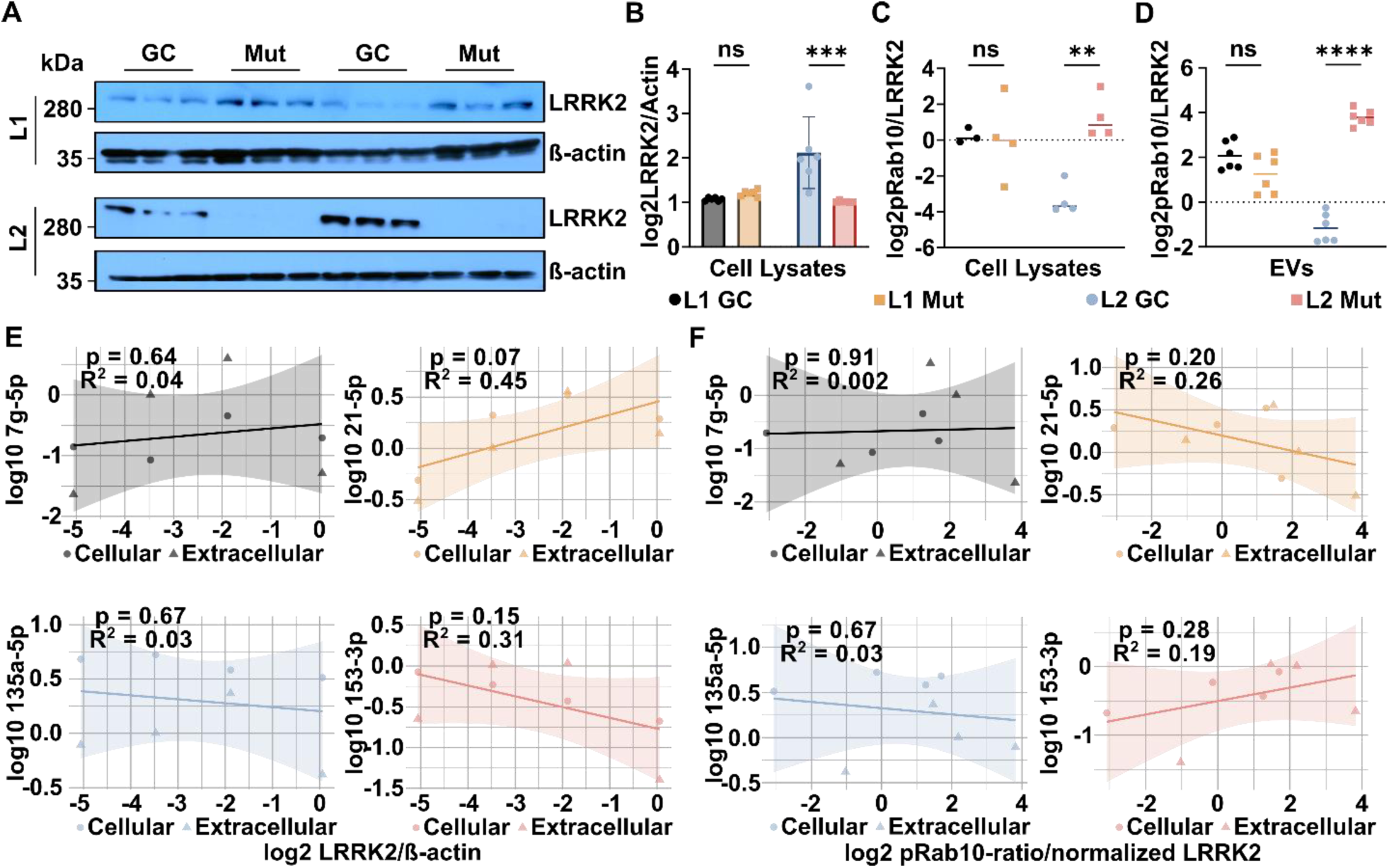
Changes of miRNA-expression values seem mostly independent of LRRK2 expression or kinase activity. **(A)** Raw western blot images displaying LRRK2 and β-Actin across both genotypes and patient lines. **(B)** After performing statistical analysis on the log-transformed LRRK2/β-Actin ratios, LRRK2 expression levels were similar in the L1 lines. In contrast, L2 Mut exhibited a significant downregulation of LRRK2 expression compared to L2 GC. **(C)** We quantified and calculated pRab10 over total Rab10 ratios in both cell lysates and **(D)** EVs and compared those to the LRRK2 ratios measured in cell lysates. Each dot represents a separate batch of cell lysates or EVs. Interestingly, kinase activity did not differ significantly in the L1 lines while L2 Mut showed a substantial increase in kinase activity in both cell lysates and EVs. **(E)** Finally, we conducted Pearson’s correlation analysis to assess the relationship between LRRK2 expression levels or **(F)** pRab10/LRRK2 ratios and the expression levels of the four validated miRNAs from our libraries. Each plot shows data from both cellular and extracellular RNA and from both lines. Each data point represents a technical replicate, with dots indicating cellular RNA data and triangles representing extracellular RNA data. Except for the correlation between miR-21-5p and LRRK2 expression, which showed a strong trend towards significance (p = 0.07), variations in LRRK2 expression or activity did not appear to linearly influence miRNA expression levels.

As a next step, we quantified pRab10/Rab ratios in cell lysates and EVs as a proxy of LRRK2 activity. Calculated pRab10/Rab10 ratios were normalized to LRRK2 expression levels. In cell lysates, the main effect of patient cell line was not statistically significant (F(1,11) = 2.46, p = 0.145). The main effect of mutation status was significant (F(1,11) = 10.37, p = 0.008). Furthermore, a significant interaction between the two factors was found (F(1,11) = 10.22, p = 0.009). After performing Tukey’s test for multiple comparisons, the difference of the pRab10/Rab10 ratios was not significant in the L1 lines (p = 0.99), while it was statistically significant in the L2 lines (p = 0.004) (Figure 5C). In EVs, there was no significant main effect of the patient cell line on the pRab10/Rab10 ratios (F(1,20) = 1.75, p = 0.201). However, there was a significant main effect of mutation status (F(1,20) = 60.09, p<0.001), indicating that mutation status had a strong influence on the pRab10/Rab10 ratios. Additionally, a significant interaction effect was observed between the two factors (F(1,20) = 116.14,p<0.001). After performing Tukey’s test for multiple comparisons, analog to the findings in cell lysates, ratios were not different in L1 lines (p = 0.17), while being significant in the L2 lines (p<0.0001) (Figure 5D).

Finally, we performed a correlation analysis between the normalized expression levels of miRNAs 7g-5p, 21-5p, 135a-5p, and 153-5p and either LRRK2 protein expression (Figure 5E) or pRab10/Rab10 ratios (Figure 5F). While no statistically significant correlations were identified, the association between miR-21-5p expression and LRRK2 protein levels showed a notable trend toward significance (R² = 0.45, p = 0.07).

## Discussion

Cell-free miRNAs were shown to be differentially expressed in patients with sporadic Parkinson’s disease across both brain tissue and body fluids, underscoring their potential as promising biomarkers [43]. While approximately 15% of PD cases are attributed mainly to specific genetic factors, studies investigating miRNA alterations in the context of fPD remain limited. In this proof-of-concept study, we focused on identifying miRNA-based biomarkers associated with the LRRK2 G2019S mutation using an iPSC-derived in vitro model for PD. Our findings demonstrate that changes in the expression of cell-free miRNAs closely mirror those observed within the intracellular miRNAome. We further identified four miRNAs consistently dysregulated in both cell-free and cellular RNA in vitro, namely miR-21-5p, miR-135a-5p and let-7g-5p. Finally, we found that the observed changes in miRNA expression did not follow a linear relationship with LRRK2 expression or activity.

We used iPSC-derived dopaminergic neurons carrying the LRRK2 G2019S mutation as an in vitro model for fPD. We confirmed the neuronal identity and maturity of the cells via the quantification of the neuronal marker MAP2. Dopaminergic cells were detected using TH. The cells displayed neuronal morphologies and stable expression of MAP2 and TH. Additionally, quantification of the transcription factor FOXA2, which is a marker for midbrain neurons [44] and plays a role in the survival of dopaminergic cells [45], showed high expression levels in all four lines compared to iPSCs. A more thorough characterization of the iPSCs used in this study, including stainings of LRRK2, quantification of additional dopaminergic markers, and dopamine uptake assays, has been reported previously [26]. We used cell culture EVs as a source for cell-free RNA, and their presence was confirmed via NTA, electron microscopy, and blotting of EV markers Alix, Flotillin-1, and CD81 [41].

In our small-RNA libraries, we found a total of 56 differentially expressed miRNAs. Notably, there was no overlap between the two patient-derived cell lines. As we were interested in identifying a LRRK2- related miRNA biomarker, we quantified the expression levels of LRRK2 using qPCR and Western Blotting. Surprisingly, there were significant differences between the L1 and L2 lines, with LRRK2 being upregulated in L1 Mut and downregulated in L2 Mut. Although the regulation of LRRK2 expression is not yet completely understood, changes in expression levels have been previously reported. For instance, LRRK2 was upregulated in colonic tissue from PD patients [46] and shown to be influenced by other PD-related proteins, such as PINK1 [47]. As LRRK2 is known to influence the cellular miRNAome [22], we anticipated the differential LRRK2 expression levels to have a major impact on our results. Remarkably, in the present study, we found that the L1 and L2 lines frequently exhibited very different miRNA profiles.

In fact, of the four miRNAs validated from our libraries, only one miRNA displayed upregulation in EVs in both cell lines, namely 135a-5p. This miRNA was previously found to be upregulated in the substantia nigra of male PD brains [48] and has been linked to dysfunctional insulin signaling [49]. Interestingly, disruptions in the insulin-signaling pathways have been suggested to play a role in both ageing and neurodegeneration [50]. One potential explanation for miR-135a-5p being dysregulated in the same direction in both cell lines despite the differences in LRRK2 expression could therefore be, that it is further downstream of LRRK2 and more reflective of a general PD pathology, in this case triggered by an interplay of the LRRK2 G2019S mutation and other factors, such as the genetic background.

The other three miRNAs validated from our libraries exhibited opposing behaviors between the two cell lines. Among those, let-7g-5p was consistently upregulated in vitro in L1, whereas it was downregulated in L2. It is regarded as a protective miRNA due to its demonstrated ability to repress α-synuclein protein levels [51], and its potential role in disease-modifying therapies is under active investigation [52]. Additionally, increased let-7 levels have been reported to mitigate the pathogenic effects associated with LRRK2 [22]. The differing patterns of let-7g-5p dysregulation observed in vitro may therefore reflect varying degrees of compensatory response mechanisms aimed at counteracting LRRK2-driven pathology [53].

miR-21-5p was consistently upregulated in hDaNs of L1, whereas it was downregulated in L2. MiR- 21 has been linked to various central nervous system disorders, including Alzheimer’s disease and PD, and is known to regulate inflammatory processes and apoptosis [54]. Furthermore, upregulated levels of miR- 21 were shown to increase expression levels of α-synuclein [55] and in the substantia nigra of six PD brain samples [56]. The altered expression levels of miR-21 may therefore reflect dysregulations in α-synuclein expression or disruptions within the broader regulatory network influenced by miR-21.

miR-153, which was found to be upregulated in both cellular and cell-free RNA from the L2 lines, has previously been associated with PD. Multiple studies have demonstrated that miR-153, alongside miR- 7, plays a role in suppressing the accumulation of α-synuclein [57; 58]. Recently, we reported reduced levels of miR-153-3p, a mature form derived from the 3’ arm of miR-153, in the plasma of LRRK2 mutation carriers [59]. The increased levels of miR-153 observed in the L2 line may therefore represent a compensatory mechanism in response to the altered LRRK2 activity in this line.

Although other factors may contribute to the opposing miRNA dysregulation patterns observed in the two lines, we strongly believe that LRRK2 plays an important role as it is a known critical master regulator influencing the miRNAome [23]. Importantly, our data did not reveal any straightforward linear correlation between LRRK2 expression levels or activity and miRNA alterations. This lack of linearity underscores the possibility that miRNA changes might be influenced by LRRK2 in an indirect or context-specific manner, rather than through a simple dose-dependent relationship. It is also plausible that miRNA dynamics are influenced by compensatory mechanisms, feedback loops, or other post-transcriptional regulatory factors that obscured a direct association with LRRK2 in our experimental setup.

On a final note, we found a statistically significant correlation between changes in the cellular and cell-free miRNAome in our in vitro model. This data suggests that changes in the cellular miRNAome that might occur as a response to PD specific stressors are well reflected in the cell-free miRNAome.

### Limitations

The protocols for generating small RNA libraries differed slightly between the two patients due to variations in the time points for collecting cell culture supernatants. Since the libraries were not generated simultaneously, we cannot fully exclude the possibility of batch effects. However, the validation of dysregulated miRNAs identified in the libraries was performed using an independent batch of cell-free RNA, where only supernatants collected between days 14 and 23 were used. Experiments including the inhibition of LRRK2 could have further elucidated the relation between LRRK2 activity and changes in the miRNAome. Lastly, a critical consideration in this study was how to normalize RT-qPCR data for miRNA expression levels. Unlike cellular mRNA experiments, where well-established housekeeping genes such as GAPDH are commonly used for normalization, a definitive gold standard for miRNA quantification is still lacking. Therefore, we utilized miR-16 as an internal control, as it has been used in several prior studies, including those related to neurodegeneration [60].

### Conclusions

let-7g-5p, miR-135a, and miR-21-5p appear to be promising targets for future biomarker studies. Further studies will be necessary to validate the identified miRNA targets due to the limited sample size in this study. Our findings suggest that identical disease-causing mutations can result in distinct alterations in miRNA patterns, potentially reflecting differences in genetic background or compensatory cellular mechanisms. Characterizing the cellular miRNAome of patient-derived hDaNs could be employed in the future to single out promising miRNAs for each patient and to identify and monitor individually altered molecular pathways. An alternative approach could involve analyzing miRNA profiles, rather than relying solely on individual miRNA readouts, to gain a more comprehensive understanding of the regulatory landscape.

### Declaration of generative AI and AI-assisted technologies in the writing process

During the preparation of this work the authors used ChatGPT to check and increase readability. After using this tool, the authors reviewed and edited the content as needed and take full responsibility for the content of the published article.

---

**Supplemental Figure 1.**
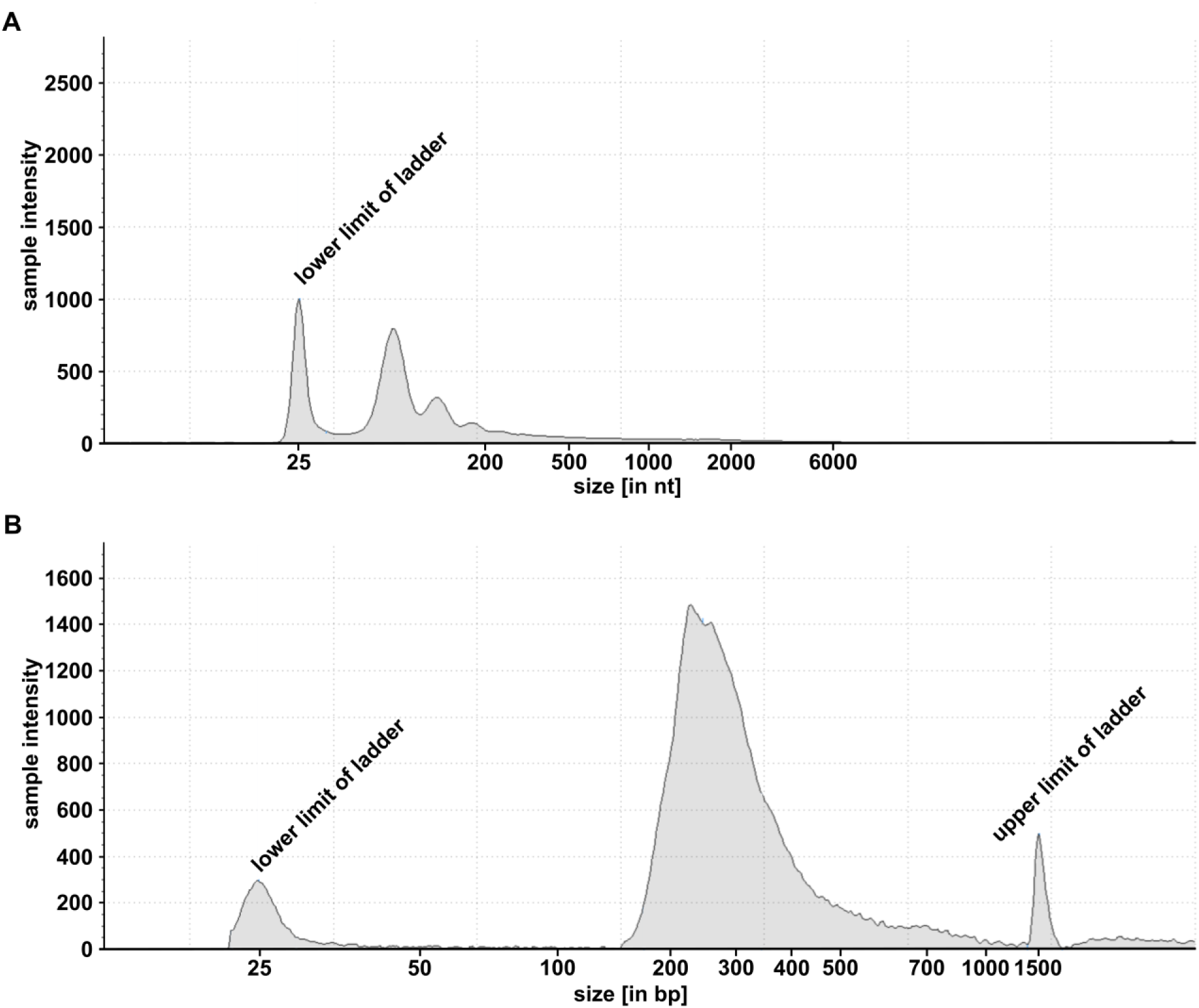
Representative electropherograms of RNA input and library. **(A)** Electropherogram of cell-free RNA isolated from EVs derived from L2 Mut. Notably, no peak at 5000nt was observable, indicating the absence of rRNA. **(B)** Electropherogram of one of the L2 Mut libraries after the second bead purification. Sequencing of the libraries was done at a higher length to allow the inclusion and better mapping of potentially present mRNAs, which is not part of the present study.

**Supplemental Figure 2.**
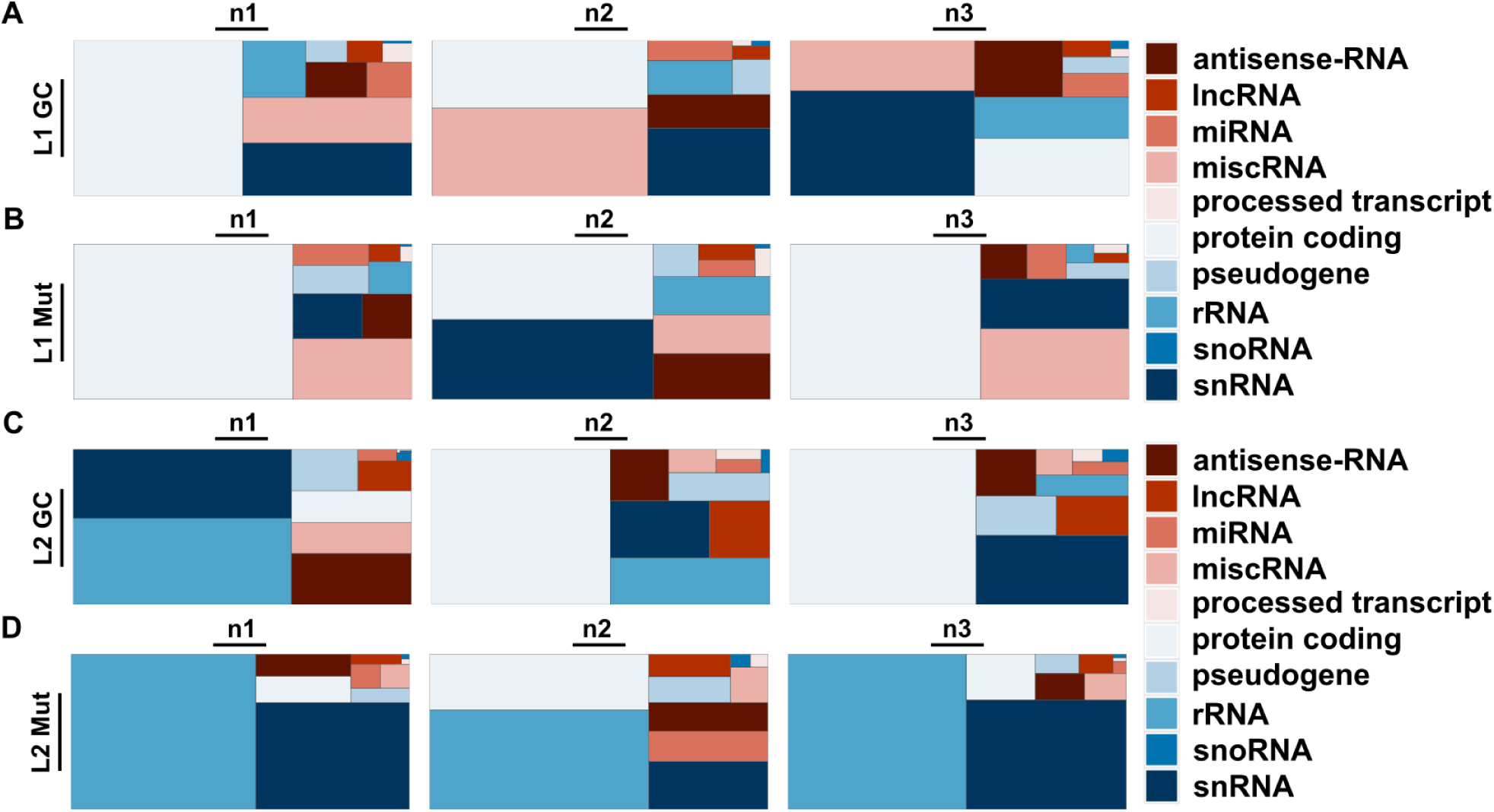
Overview of biotype composition in the generated libraries. Size of the square represents the abundance of a given RNA biotype with higher abundance resulting in larger squares. Only the ten most abundant RNA biotypes are shown. The three boxes per row (n1-n3) represent the independent differentiations that were used to generate the libraries **(A)** Libraries generated from material from L1 GC, **(B)** L1 Mut, **(C)** L2 GC and **(D)** L2 Mut.

**Supplemental Figure 3.**
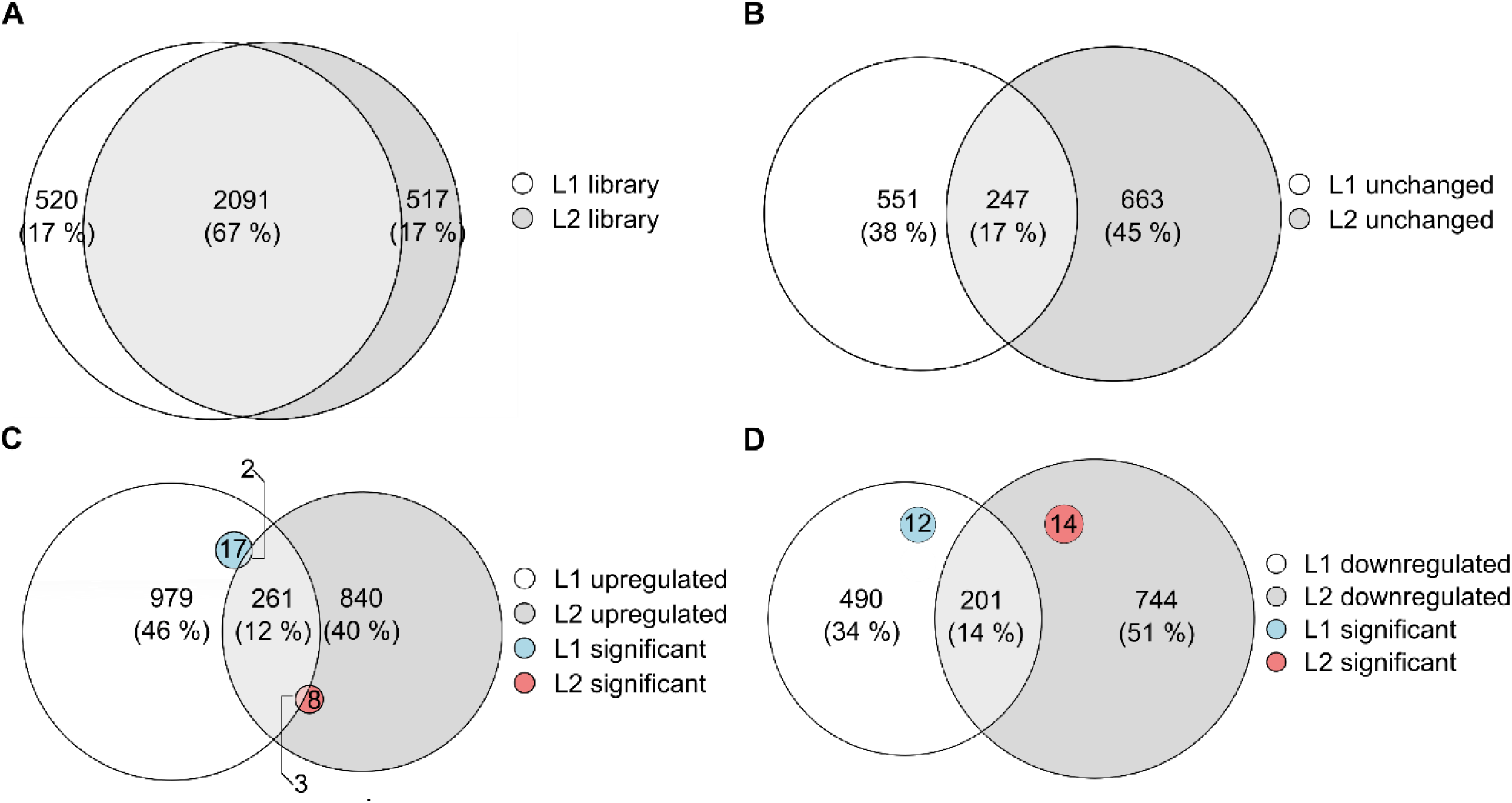
Overview of overlaps between the L1 and L2 libraries. **(A)** A total of 2091 miRNAs were found to be present in both libraries **(B)** Of the miRNAs identified in the libraries, 798 (L1 Mut) and 910 (L2 Mut) did not show differential expression levels. Of these unchanged miRNAs, 247 were found in both patient lines. **(C)** Of the miRNAs found to be upregulated, 256 were shared by L1 Mut and L2 Mut. While there was no overlap after correction for multiple testing, 5 of the significantly upregulated miRNAs were among the 261 miRNAs that trended towards upregulation. **(D)** In L1 Mut, 703 miRNAs were downregulated compared to 959 in L2 Mut. Between them, 201 miRNAs were downregulated in both patient lines. After correction for multiple testing, no overlap between the lines was found.

